# Microbiota differences of the comb jelly *Mnemiopsis leidyi* in native and invasive sub-populations

**DOI:** 10.1101/601419

**Authors:** Cornelia Jaspers, Nancy Weiland-Bräuer, Martin Fischer, Sven Künzel, Ruth A. Schmitz, Thorsten B.H. Reusch

## Abstract

The translocation of non-indigenous species around the world, especially in marine systems, is a matter of concern for biodiversity conservation and ecosystem functioning. While specific traits are often recognized to influence establishment success of non-indigenous species, the impact of the associated microbial community for the fitness, performance and invasion success of basal marine metazoans remains vastly unknown. In this study we compared the microbiota community composition of the invasive ctenophore *Mnemiopsis leidyi* in different native and invasive sub-populations along with characterization of the genetic structure of the host. By 16S rRNA gene amplicon sequencing we showed that the sister group to all metazoans, namely ctenophores, harbored a distinct microbiota on the animal host, which significantly differed across two major tissues, namely epidermis and gastrodermis. Additionally, we identified significant differences between native and invasive sub-populations of *M. leidyi*, which indicate, that the microbiota community is likely influenced by the genotypic background of the ctenophore. To test the hypothesis that the microbiota is genotypically selected for by the ctenophore host, experiments under controlled environments are required.

## INTRODUCTION

The multicellular organism and its associated microbiota, commonly named metaorganism, is attaining large public and scientific attention. For model organisms it has been shown that the microbiota contributes to host fitness and can be essential for host functioning (Bosch et al., 2011; Rook et al., 2017; Jaspers et al., 2019). Ctenophores, being a basal metazoan phylum (Dunn et al., 2008; Whelan et al., 2017), have not been considered in the metaorganism context up to now (Bosch et al., 2011). However, it is expected that a more detailed understanding about host-microbe interactions in this basal metazoan can reveal important information about metaorganism evolution. For example, in other marine basal metazoans, it has been shown that bacteria have a crucial impact on life history decision such as metamorphose and settlement induction to the benthos (as reviewed in Wahl et al., 2012). Additionally, bacterial exudates have been documented to trigger sexual reproduction in chanoflagellates (Woznica et al., 2017) as well as the development of multicellularity (Alegado et al., 2012). Hence, a more detailed understanding about host-microbe interaction at the base of the metazoan tree of life is essential to understand evolutionary trajectories of the metaorganism.

The ctenophore *Mnemiopsis leidyi* is not only a basal metazoan species, but can be regarded as an ecological model organism (Jaspers et al., 2018). This is attributed to the fact that *M. leidyi* has a long invasion history in western Eurasia (as reviewed in Jaspers et al., 2018), with large and widespread impacts on invaded ecosystems (e.g. Kideys 2002; Tiselius et al., 2017). For western Eurasia, two distinct invasion events can be differentiated with a first introduction into the Black Sea which subsequently spreading throughout adjacent waters including the Mediterranean Sea, and a more recent introduction into north-western Europe (Reusch, et al., 2010; Jaspers et al., 2018). Both introductions can be traced to different sub-population origin in its native habitat with the north European invasion stemming from the NE coast of the USA, while the southern invasion can be traced back to the Gulf of Mexico and SE coast of the USA (Bolte et al., 2013; Bayha et al., 2015).

Some earlier studies characterized the associated microbiome of *M. leidyi* in invaded (Hao et al., 2015) and native (Daniels et al., 2012) populations but came to inconclusive results. For example, the study by Hamman et al. (2015) claimed that *M. leidyi* epithelia in invaded habitats do not contain an associated microbiome (Hammann et al., 2015), which was, however, not confirmed by others (Dinasquet et al., 2012). The aim of the current study was to characterize the microbiome of the ctenophore *Mnemiopsis leidyi* in relation to the host genotype and considering different host tissues using standardized sampling and sequencing methods. These data provide a first indication on the stability and evolution of the microbiome through time during invasions into novel habitats.

## MATERIALS AND METHODS

### Sampling of ctenophores

*Mnemiopsis leidyi* were sampled in northern and southern areas of its native as well as invasive distribution range. Namely, samples originated from Mosquito Lagoon, Florida, SE USA, Atlantic Coast; Woods Hole, Cape Cod, NE USA, Atlantic Coast; Marseille, France, NE Mediterranean Sea and Helgoland, Germany, German Bight, North Sea. Animal were caught by use of a dip net, individually kept in sterile filtered sea water for up to 6 hours before samples were generated. All animals were ensured to have empty guts upon dissection. Tissue samples were attained by cutting the animals with a sterile scalpel on sterile petri dishes and samples immediately frozen at −80°C until processing. All samples were attained during summer 2012.

### Nucleic acid isolation and 16S rRNA gene amplicon sequencing

DNA was sampled from two different tissue types of *M. leidyi*, namely mucus from the outer epithelia (mesoglea: 1 mL v/v) as well as from the gastric cavity (pharynx: 200 μL v/v). Additionally, surrounding (inoculum) water samples (5 to 11 replicates per site) were taken and consisted each of 1 L pre-filtered (10μm isopore membrane filter to remove larger zooplankton – Millipore Merck, Darmstadt, Germany, TCTP 04700) seawater concentrated on a 0.22 μm pore size membrane filter (Millipore Merck, Darmstadt, Germany, GPWP 04700). DNA was isolated using the Wizard Genomic DNA Purification Kit (Promega, Madison, USA). DNA isolation was performed for at least 5 replicates of each sample type. PCR amplification of the V1–V2 hypervariable region of the 16S rRNA gene was performed with primers V2_A_Pyro_27F (5′-CGTATCGCCTCCCTCGCGCCATCAGTCAGAGTTTGATCCTGGCTCAG-3′) and V2_B_Pyro_27F (5′-CTATGCGCCTTGCCAGCCCGCTCAGTCAGAGTTTGATCCTGGCTCAG-3′). Amplicons were size checked and purified using MinElute Gel Extraction Kit (Qiagen, Hilden, Germany). Purified amplicon DNAs were quantified using Quant-iT picoGreen kit (Thermo Fisher Scientific, Waltham, USA) and equal amounts of purified PCR products were pooled for subsequent sequencing using the Illumina MiSeq platform with library preparation according to manufacturer’s instructions at Max-planck Institute for Evolutionary Biology.

### Data processing and bioinformatics

All steps of sequence processing were conducted with the program mothur v1.27.0 (Schloss et al., 2009). The Greengenes reference taxonomy (DeSantis et al., 2006) available on http://www.mothur.org/wiki/Greengenes-formatted_databases was used with reference sequences trimmed to the V1-V2 primer region to improve accuracy of classification (Werner et al., 2012). The sequence depth of all 117 analyzed samples ranged from 23,622 to 675,908 single sequences. The aligned dataset was used to compute a distance matrix (dist.seqs) for binning sequences into operational taxonomic units (OTU) by average neighbor clustering (cluster.split). 2829 OTUs at a 97 % similarity threshold (roughly corresponding to species level) distinction were considered. All downstream computations were performed in R v2.15.1 (R_Core_Team 2012), following published protocols (Weiland-Bräuer et al., 2015).

### Genotyping

Genomic DNA extracts from *M. leidyi* epithelia tissue samples were used for genotype analyses of all sub-populations using 7 highly polymorphic microsatellite markers (Reusch et al., 2010). Sequencing analyses were conducted on an ABI 3130 genetic analyzer using the internal size standard Rox-350 (Applied Biosystems). Allele sizes were scored using the software GENEMARKER v1.91 (SoftGenetics, LLC) with all sample processing and statistical analyses conducted as outlined previously (Reusch et al., 2010; Jaspers et al., 2018). Pairwise F_ST_ values were computed in ARLEQUIN v3.5.1.2 (Excoffier et al., 2010), using non-parametric permutation procedures. All tests were performed with an initial alpha level of 0.05 and corrected following the Benjamini-Hochberg False Discovery Rate (B-H FDR) procedure (Benjamini et al., 2001). The software STRUCTURE v2.3.4 (Pritchard et al., 2010) was used to infer genetic clustering without a priori assumption about expected number of clusters. Probabilities were calculated for k ranging from 1 to 4 with five replicates for each k and the most likely number of k’s inferred through the Evanno method (Evanno et al., 2005) using the internet interphase STRUCTURE HARVESTER (Earl et al., 2012).

## RESULTS

### Phylogenetic composition of associated bacteria reveals epithelial and sub-population – specific colonization of the ctenophore

The microbiota associated with different epithelia and sub-population of *M. leidyi* as well as the bacterial communities of ambient water were analyzed by Illumina sequencing the V1-V2 region of the 16S rRNA gene. DNA isolation, PCR amplification, and sequencing was successful with almost all samples resulting in sufficient numbers of replicates for each sample (n = 5 up to 17). The microbiota associated with epidermis and gastrodermis of two native and two invasive *M. leidyi* sub-populations were analyzed in comparison to the respective ambient water. Sequence analysis of 117 samples revealed sequence depths of 23,622 to 675,908 single sequences. In total, 2,829 OTUs at a 97 % similarity threshold (roughly corresponding to species level) were considered. Abundances of OTUs (97 % sequence similarity) between individual samples are depicted in Fig. 1 at genus level. Main representative bacterial classes in the samples were Alphaproteobacteria, Gammaproteobacteria, Mollicutes, Cyanobacteria and Bacteroidia. While variability of OTU abundances between the different sample types was significant, in the following the different comparisons are considered in more detail.

**Figure 1:**
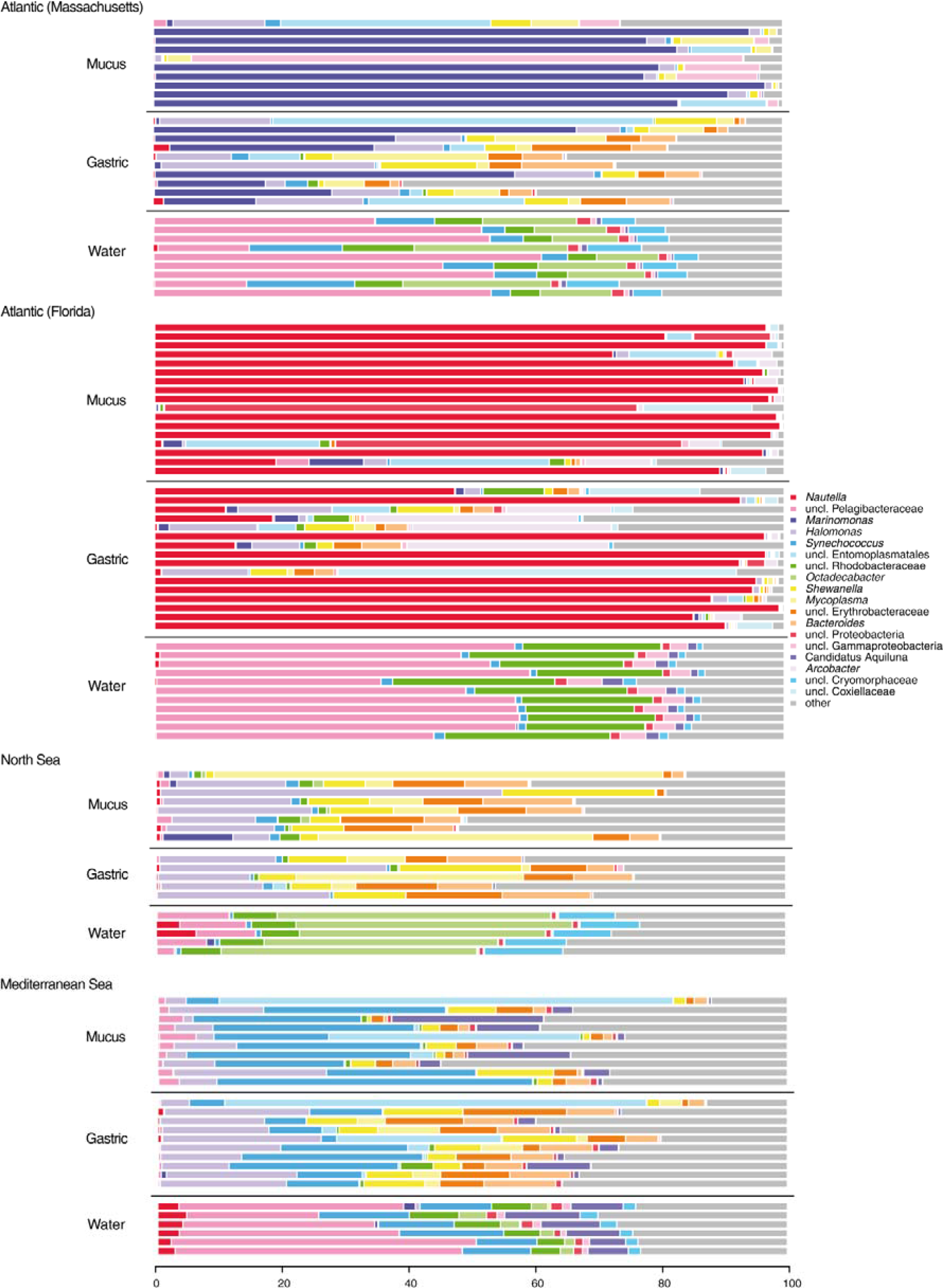
Bacterial composition patterns of individual *M. leidyi* samples. Microbial communities associated with different epithelia of several sub-populations (native and invasive) of *M. leidyi* were analyzed by sequencing the V1-V2 region of 16S bacterial rRNA gene. Taxonomic assemblages inferred from non-chimeric operational taxonomic units using mothur at a 97 % pairwise similarity cut-off; distribution patterns were calculated using R. OTU abundances were summarized at the genus level. Bar plots are grouped according to the sample type characteristic “provenance”, splitted into “mucus”, “gastric” and “water” each including at least 5 replicates. Taxa with a mean relative abundance of < 1 % across all samples are subsumed under “other”.

### Significant difference between bacterial communities from ctenophores and water columns

*First*, beta diversity analysis demonstrated that the bacterial community of the ctenophore was always significantly different from the community present in the water column (Fig. 2, Tab. 1). Water samples were mainly dominated by uncultured Pelagibacteraceae and uncultured Rhodobacteraceae of class Alphaproteobacteria, and *Synechoccus* of class Cyanobacteria, whereas the ctenophores were mostly colonized by *Halomonas*, *Shewanella*, and *Marinomonas* of Gammaproteobacteria, and Mollicutes.

**Figure 2:**
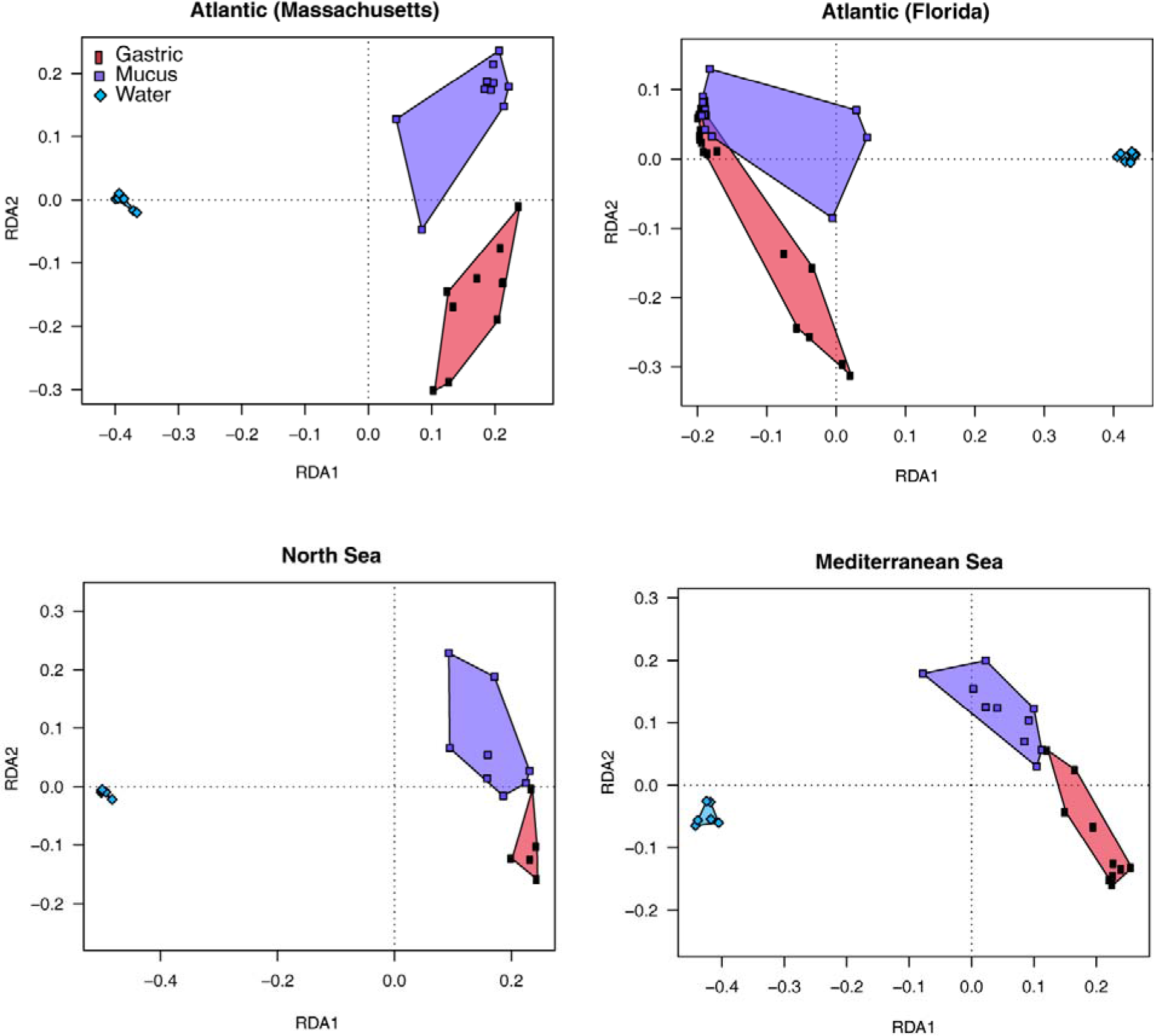
Redundancy analysis (RDA) models of Hellinger-transformed OTU abundances comparing *M. leidyi* associated bacteria of the gastric cavity and the mucus with the respective ambient water. Graphical representation (distance plots) of redundancy analysis (RDA) model of Hellinger-transformed OTU abundances. Each point represents the whole microbial community of an individual sample from gastric cavity, mucus and ambient water of the respective sub-population. Groups of related sample points are framed by polygons filled with a corresponding color to elucidate distribution and variability of sample groups in the ordination space.

**Table 1:** Results of pairwise tests in beta diversity analysis. Tests were conducted for specified comparisons. Significant results (alpha = 0.05) are highlighted (boldface).

| Comparison | *P*-value |
| --- | --- |
| Atlantic (Massachusetts): Gastric - Mucus | **0.004** |
| Atlantic (Massachusetts): Gastric - Water | **0.001** |
| Atlantic (Massachusetts): Mucus - Water | **0.001** |
| Atlantic (Florida): Gastric - Mucus | **0.030** |
| Atlantic (Florida): Gastric - Water | **0.001** |
| Atlantic (Florida): Mucus - Water | **0.001** |
| North Sea: Gastric - Mucus | 0.059 |
| North Sea: Gastric - Water | **0.001** |
| North Sea: Mucus - Water | **0.002** |
| Mediterranean Sea: Gastric - Mucus | **0.002** |
| Mediterranean Sea: Gastric - Water | **0.001** |
| Mediterranean Sea: Mucus - Water | **0.001** |
| Atlantic (Massachusetts) - Atlantic (Florida) | **0.001** |
| Atlantic (Massachusetts) - North Sea | **0.001** |
| Atlantic (Massachusetts) - Mediterranean Sea | **0.001** |
| Atlantic (Florida) - North Sea | **0.001** |
| Atlantic (Florida) - Mediterranean Sea | **0.001** |
| North Sea - Mediterranean Sea | **0.001** |

### Tissue variation of ctenophore-associated bacteria

*Se*cond, sequencing results based on the 16S rRNA gene amplicon sequencing revealed differences in the bacterial composition among two different tissues, the outer mucus layer on the epidermis, and the epithelium in the gastric cavity. For all populations the tissues harbored significantly specific microbiota (*P* = 0.001), except for the North Sea where microbiota composition differed marginally (*P* = 0.059). Mucus from native Atlantic (Massachusetts) animals was dominated by *Marinomonas* and unclassified Pelagibacteraceae, whereas their gastric cavity was mainly colonized by *Shewanella*, *Mycoplasma*, *Bacteroides*, unclassified Erythrobacteraceae and a multitude of low abundant OTUs. Both, mucus and gastric cavity from native Atlantic (Florida) animals were dominated by *Nautella*; mucus additionally comprised unclassified Entoplasmatales, whereas gastric cavity was additionally colonized by unclassified Erythobacteracaeae, *Bacteroides* and *Arcobacter*. Invasive Mediterranean mucus and gastric samples both, mostly comprised *Synecchococcus*, mucus samples additionally contained unclassified Entoplasmatales and Candidatus Aquiluna, whereas gastric samples comprised *Halomonas*, *Shewanella*, *Mycoplasma*, unclassified Erythrobacteraceae and *Bacteroides*. Main representatives in mucus as well as gastric samples of invasive North Sea animals were *Halomonas*, *Shewanella*, *Mycoplasma*, unclassified Erythobacteraceae and *Bacteroides*. Mucus and gastric cavity community patterns of those animals were almost similar, however slight differences were observed in abundances of main representatives and presence of different low abundant taxa.

### Sub-population specific community patterns of M. leidyi

Third, bacterial community analysis of native (Atlantic – Massachusetts; Atlantic - Florida) and invasive (North Sea, Mediterranean Sea) *M. leidyi* sub-populations revealed significant differences due to provenance. Similarly, population structure analyses using the polymorphism of seven microsatellite loci revealed that all tested sub-populations were significantly differentiated (F_ST_ range 0.03 to 0.33; *P*<0.05 after correction for multiple testing, Table 2). The largest observed population differentiation was found between the northern invasive sub-population (Helgoland, North Sea) and the southern native sub-population (Florida, Atlantic Coast) with F_ST_=0.333 (*P*<0.0001, Table 2), while the smallest population structure difference was observed between native northern sub-populations (Massachusetts, NE USA) in comparison with their recently introduced counterparts in Northern Europe (Helgoland, North Sea) with F_ST_=0.032 (*P*<0.05, Table 2). While the northern native and invasive sub-populations showed significant differences in the pairwise genetic distance calculation analyses, this was not confirmed by data exploration using a Bayesian clustering approach, as the STRUCTURE analyses indicated 3 clusters (Fig. 3), without resolving the northern native and invasive sub-populations (see Table 2).

**Figure 3:**
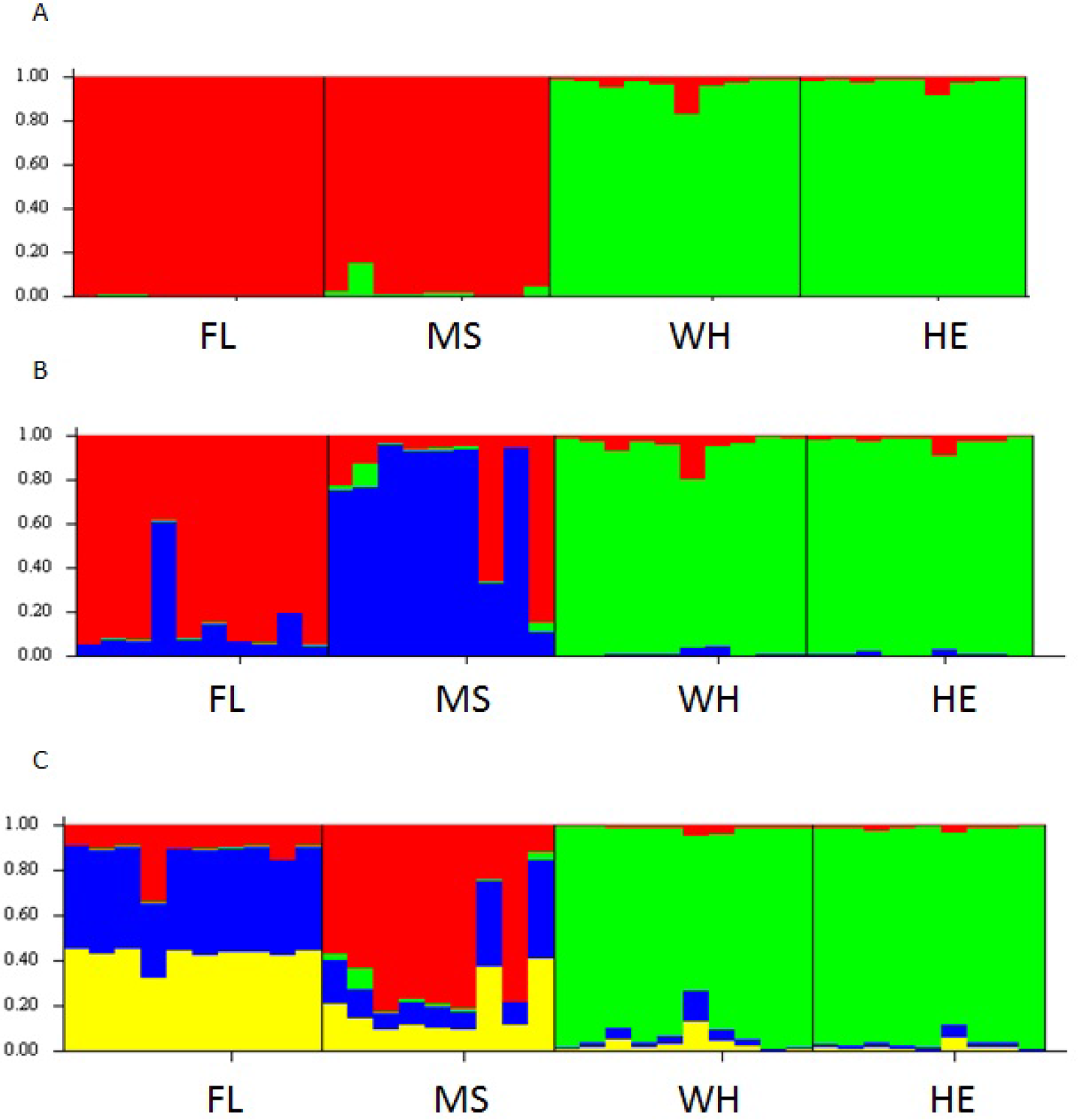
Structure plot population differentiation based on 7 highly polymorphic microsatellite data. STRUCTURE HARVESTER (Earl et al., 2012) was used to calculate the most likely number of clusters (ΔK). The resulting significant number of clusters was one. However, visual inspection of the output files created by Structure 2.3.4 (Panel A with K = 2, panel B with K = 3, and panel 4 with K = 4), clearly showed the presence of three clusters (K = 3), while with this method the 4^th^ cluster could not be resolved (but see Table 2).

**Table 2:** Pairwise comparison of genetic differentiation (F_ST_) between native and invasive *M. leidyi* sub-populations using highly polymorphic microsatellite markers. The four tested sub-populations are abbreviated with southern native = FL (Florida, USA, SE Atlantic coast), northern native = WH (Woods Hole, Massachusetts, USA, NE Atlantic coast), southern invasive = MS (Marseille, France, NE Mediterranean Sea) and northern invasive = HE (Helgoland, German Bight, North Sea). Significant comparisons are highlighted in bold with *P*-levels abbreviated by *P*<0.05*, *P*<0.001**, *P*<0.0001*** (p-values corrected for multiple testing).

|  | MS | FL | WH | He |
| --- | --- | --- | --- | --- |
| MS | - |  |  |  |
| FL | **0.078\*\*\*** | - |  |  |
| WH | **0.277\*\*\*** | **0.157\*\*\*** | - |  |
| He | **0.333\*\*\*** | **0.205\*\*\*** | **0.032\*** | - |

With regard to associated bacteria community differences, RDA plots demonstrated that every single sub-population hardboard a specific microbiota. Native sub-populations are significantly different and share just few OTUs. Invasive sub-populations shared a more similar microbiota and were comparable with native Atlantic (Massachusetts) animals, however significantly different to native Atlantic (Florida) animals (Fig. 4).

**Figure 4:**
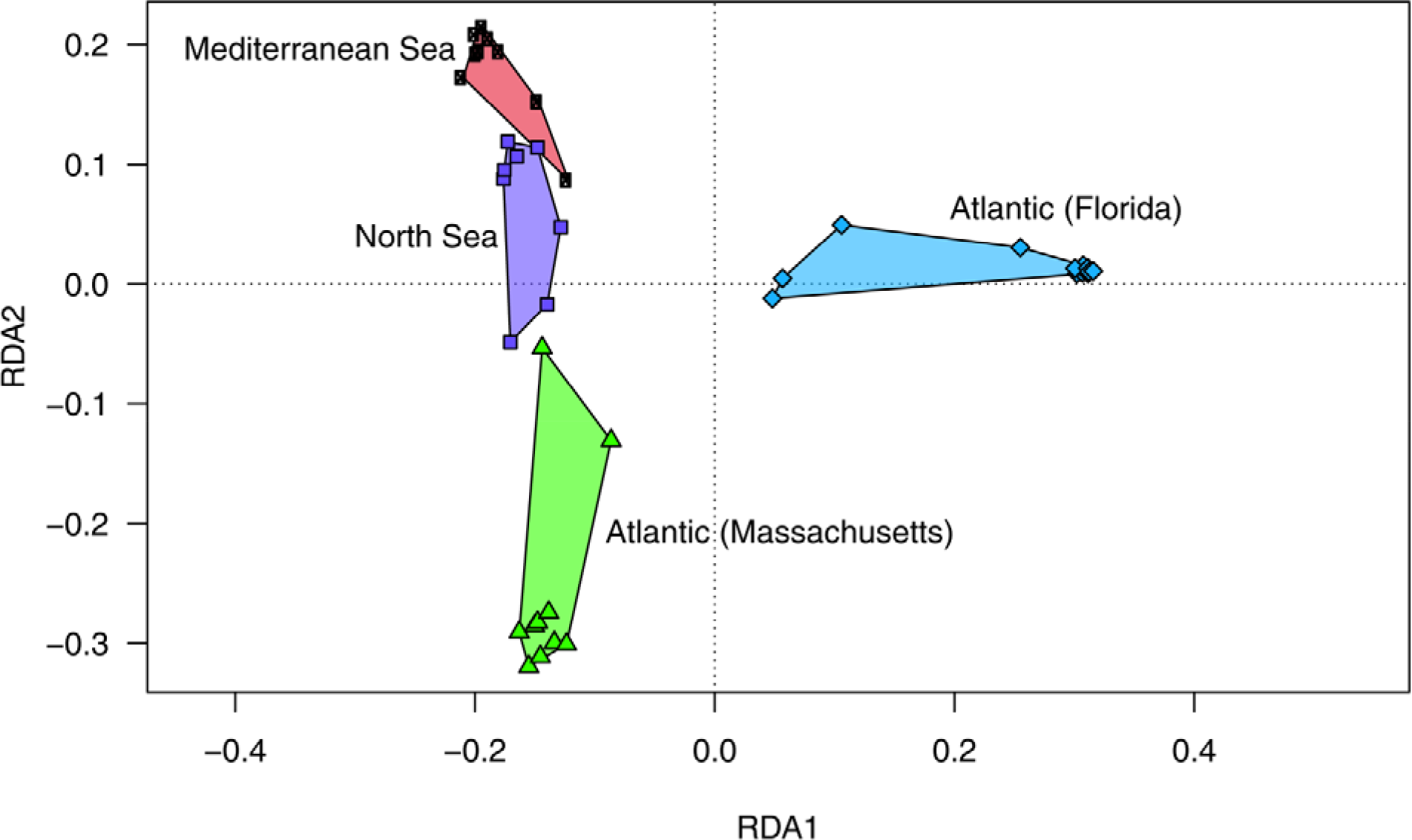
Redundancy analysis (RDA) models of Hellinger-transformed OTU abundances comparing *M. leidyi* associated bacteria of different sub-populations. Distance plots of redundancy analysis (RDA) model of Hellinger-transformed OTU abundances. Each point represents the whole microbial community of an individual sample associated with the respective native and invasive sub-population. Groups of related sample points are framed by polygons filled with a corresponding color.

## DISCUSSION

This is the first study which compared the microbiota community composition of native and invasive sub-populations of the comb jelly *Mnemiopsis leidyi* using standardized high throughput amplicon sequencing methods based on the V1V2 region of the 16S rRNA gene. We demonstrated that the microbiota community composition was significantly different among all investigated sub-populations. The microbiome was specific to each sub-population and significantly different from the respective surrounding water. Contrary to the previous report that the mucus and epidermis of *M. leidyi* is free from associated microbes (Hammann et al., 2015), we observed a diverse and distinctly different microbial community composition on epithelia tissue as well as the gastrodermis, in line with Dinasquet et al. (2012).

While a core microbiome has been shown to remain throughout development in some scyphozoan jellyfish species (Lee et al., 2018), the microbiome of the scyphozoan jellyfish *Aurelia aurita* showed significant life-stage dependent differences in the microbial species community composition (Weiland-Bräuer et al., 2015). Ctenophores are holoplanktonic throughout their life cycle and do not have a benthic polyp stage, hence probably less likely to change microbiome throughout development. For this study, we have sampled only adult *M. leidyi*, thereby excluding any potential life-stage dependent differences, which cannot be excluded for other studies as an indication of life stage is often missing. In invasive habitats, the microbiome of *M. leidyi* has been compared to other native ctenophore species, which indicated a significant difference between the microbiota community composition of all studied species (Hao et al., 2015). Irrespectively of differences in the bacterial species community composition, all investigated *M. leidyi* microbiomes were dominated by Proteobacteria, though variable through time (Hao et al., 2015). A change in bacteria species community composition with season has similarly been observed in southern native and northern invasive habitat of *M. leidyi* (Daniels et al., 2012; Hao et al., 2015). We analysed *M. leidyi* from different regions in native and invasive habitats but during the same seasonal time point. Hence, we do not expect that our samples were biased by the seasonal variation observed in other studies.

Multivariate analyses indicate that invasive and native northern sub-populations are more similar compared with southern native and invasive sub-populations. For northern native sub-populations, *Marinomonas* spp. dominated the bacteria community. However, for invasive northern sub-populations, they were only observed with a limited contribution to the microbiome during 2012. This is contrary to previous findings where during 2009/2010 *Marinomonas* spp. had been documented to be the most important bacteria component in *M. leidyi* from the invasive northern sub-population i.e. Helgoland Road (Hao et al., 2015) and the Gulmar Fjord (Dinasquet et al., 2012). A change in the bacteria community composition with reduced contribution of *Marinomonas* spp. could be related to a significant change in winter conditions as observed during the early 2010’s, where a series of cold winters had led to a retraction of the range expansion of *M. leidyi* in northern Europe with the extinction of *M. leidyi* in e.g. the Gulmar Fjord (Jaspers et al., 2018). Hence, drastic physical conditions and extinction of local populations could be reflected in a change of the microbiome but deserve further investigation. Native southern sub-populations showed a large difference in the microbiota community composition compared to the invasive southern sub-populations. Specifically, the native southern microbiome of *M. leidyi* was dominated by *Nautella* (Alphaproteobacteria), while this genus was absent from invasive *M. leidyi*. In line with this, southern invasive and native *M. leidyi* sub-populations have also been differentiated for the longest time (Jaspers et al. 2018), which might explain their difference in microbiome community composition. Similar to our results, in a previous study, Alphaproteobacteria had been shown to be key bacteria community members of *M. leidyi* from the native southern range in the Gulf of Mexico during late spring and summer (Daniels et al., 2012). For the Mediterranean Sea, no previous microbiome characterization has been conducted. We observed a large fraction of the Cyanobacteria *Synechococcus* in mucus and gastric samples. Even though this is a free living bacterium with a widespread distribution in marine and freshwater systems and could be a contamination from the sample preparation, Cyanobacteria have been also shown to contribute to the core microbiome of native southern *M. leidyi* during certain periods (Daniels et al., 2012).

Genotype impact on microbiota community composition has been investigated in polyp colonies from three different distinct sub-population of the jellyfish *Aurelia aurita* in northern Europe, cultured for ten years under similar conditions. Weiland-Bräuer et al. (2015) found that the microbiota community composition showed significant differences, indicating that jellyfish genotype had an impact on the microbiota community composition. We found the lowest population differentiation between native NE Atlantic sub-population (Massachusetts) and the invasive northern sub-population (Helgoland, North Sea). These population structure results are in agreement with molecular results (Reusch et al., 2010) and a recent investigation summarizing the range expansion of *M. leidyi* in western Eurasia, where the northern introduction has been shown to be very recent, dating back to 2005 (Jaspers et al., 2018). Irrespectively, we show that invasive sub-populations in general share a more similar microbiota, which is most comparable with the northern native sub-population from Massachusetts.

In conclusion, significant variance in microbiota community patterns and significant population differences between sub-populations suggest an impact of environment along with genetic background (population affiliation) on the microbiota community. Long term experiments over several generations under controlled conditions are however necessary to decipher the influence of environment. We found that *M. leidyi* harbors a diverse bacterial community composition on its epithelia. The significant difference between water and associated microbiota suggests that ctenophores select the microbes in a host specific way, thereby shaping the metaorganism as a whole. How the direct interactions are between certain microbes and host fitness need to be studied in a controlled environment.

## ACKNOWLEDGEMENTS

We thank Sören Bolte for support and data collection, as well as Katharina Bading for collection of animals in the native southern range.

